# Fixel-Based Analysis reveals microstructural visual pathway changes associated with retinal remodeling in retinitis pigmentosa

**DOI:** 10.1101/2025.11.14.688435

**Authors:** Giulia Righetti, Veronica Cuevas Villanueva, Milda Reith, Lasse Wolfram, David Merle, Laura Kühlewein, Christoph Braun, Katarina Stingl, Krunoslav Stingl

## Abstract

Retinitis pigmentosa (RP) is the most common inherited retinal dystrophy, traditionally considered a photoreceptor degenerative disorder. However, the relationship between retinal degeneration and structural alterations along central visual pathways remains poorly understood. We investigated possible structural links across the retinocortical pathway in 15 genetically characterized RP patients (RHO, PDE6A, PDE6B mutations) compared with 22 healthy controls, using spectral-domain optical coherence tomography (SD-OCT) combined with diffusion-weighted MRI and fixel-based analysis (FBA).

Retinal segmentation revealed marked outer layer thinning (outer nuclear layer: −55% foveal, −77% parafoveal) with significant inner layer thickening at the fovea (inner plexiform layer +29%, inner nuclear layer +40%). Parafoveal regions showed near-complete photoreceptor loss but inner retinal thickness approaching normal values. FBA demonstrated significant fiber density reductions in optic tracts and optic radiations, without corresponding macrostructural changes in tract cross-section, suggesting early microstructural disruption preceding atrophy. Correlation analyses revealed significant associations between parafoveal inner plexiform layer thickness and fiber density in both optic tracts (r=0.856) and radiations (r=0.768), with no correlations observed at the foveal level.

These findings demonstrate complex, region-specific retinal remodeling in RP, likely reflecting Müller cell gliosis rather than mechanical compensation. The fiber density reduction without tract atrophy suggests trans-synaptic degeneration along visual pathways. This multimodal framework provides insights into structural propagation in inherited retinal dystrophies and may serve as a biomarker for evaluating therapeutic interventions aimed at photoreceptor restoration.

## Introduction

Among inherited retinal dystrophies (IRD), retinitis pigmentosa (RP) is the most common, affecting approximately 1 in 4,000 people worldwide and representing the leading genetic cause of blindness ^1^. RP is a highly heterogeneous condition, but it is generally characterized by the progressive degeneration of photoreceptor. The disease typically begins with the rod cells death in the peripheral retina, leading to early symptoms such as night blindness (nyctalopia) and difficulties adapting to darkness. As the condition advances, cone cells in the central retina also become affected, eventually resulting in progressive central vision loss. Other reported symptoms include photopsia, characterized by the perception of flashing or flickering lights, and dyschromatopsia ^1^. In some forms of RP, referred to as syndromic RP, the retinal degeneration is accompanied by additional systemic symptoms, as seen in disorders such as Usher or Bardet–Biedl syndromes ^2^. To date, over a hundred genes have been associated with RP. Among these identified as causes of non-syndromic RP, rhodopsin (RHO) mutations affect 30-40% of autosomal dominant cases, while phosphodiesterase 6 subunit mutations (especially PDE6A, PDE6B) are only minor contributors of autosomal recessive forms ^3^. Despite this remarkable genetic diversity, the common pathway involves disruption of the phototransduction cascade, leading to photoreceptor cell death and progressive vision loss. RP subtypes linked to PDE6A, PDE6B and RHO gene variants typically exhibit an early onset of night blindness during childhood, a slow natural disease progression, and relative preservation of the inner retina and central cone function into adulthood ^1^. As all three genes encode rod-related functional proteins, these three subtypes of RP represent the typical early-onset primary rod degeneration with secondary cone loss.

Although RP has traditionally been viewed as a photoreceptor degenerative disorder, emerging evidence suggests that structural changes extend throughout the entire retinal architecture. Several studies indicate that as the outer retinal layers progressively degenerate, the inner retinal layers also become increasingly disorganized ^4-6^. Surprisingly, however, the post-receptoral retinal layers often exhibit a general thickening ^7-12^, along with a relatively longer preservation of cell numbers within the inner nuclear layer (INL) ^13^, despite the profound photoreceptors loss and the expected downstream degeneration due to reduced input. The functional implications of this relative preservation remain unclear. Inner retinal remodeling has been associated with changes in microperimetry sensitivity thresholds, implying a potential impact on visual function ^10^. Similarly, although retinal ganglion cells (RCG) appear to be structurally preserved, they exhibit abnormal spike patterns in experimental models, indicating a degree of functional instability within the retinal output pathway ^14^. A central contributor to this remodeling process appears to be the Müller glial cells, which play a key role in retinal metabolism and homeostasis. Notably, they are among the first cells to exhibit metabolic changes under retinal stress, yet their response is often incoherent and uncoordinated, leading to a loss of their structural and functional identity over time ^15^. Together, these findings highlight that RP is not solely a disorder of photoreceptor loss. Instead, the loss of photoreceptor-driven signaling triggers a cascade of remodeling events at the cellular, glial, and vascular levels ^16,17^, reshaping the entire retinal architecture in ways that are still not fully understood.

Despite extensive characterization of retinal pathology in RP, the impact of the disease on the central visual pathway structures remains poorly understood. In conditions such as glaucoma, where trans-synaptic neurodegeneration of third-order neurons (RGC) is well established, significant white matter degeneration has been observed along the entire visual pathway (for a systematic review see ^18^). Previous studies have investigated the white matter integrity of the visual pathways in RP using diffusion-weighted imaging (DWI) ^19,20^. These studies observed structural alterations of the optic nerve and a reduction of fractional anisotropy (FA) in the optic radiations compared to healthy controls. Moreover, decreased FA values have been shown to correlate with visual field loss ^19^. FA has long been used as an indirect marker of white matter integrity, reflecting the directional freedom of water diffusion within tissues ^21^. However, FA is a voxel-averaged measure and thus has limited interpretability, particularly in regions with crossing or diverging fibers, where multiple fiber populations coexist within the same voxel. To overcome these limitations, a fixel-based analysis (FBA) approach has been developed ^22^, where a “*fixel*” represents an individual fiber population within a voxel. FBA enables a more detailed characterization of white matter by accounting for both microstructural changes in fiber density (FD) and macrostructural, morphometric changes in fiber-bundle cross-section (FC), which can also be combined into a single metric of fiber density and cross-section (FDC) to estimate total intra-axonal volume changes within a fiber pathway. In recent years, FBA has been increasingly applied to investigate white matter alterations in various neurodegenerative diseases, including Alzheimer’s disease ^23,24^, multiple sclerosis ^25^, and glaucoma ^26-28^, demonstrating its potential for detecting subtle and disease-specific patterns of structural change. Applying these advanced techniques to RP patients offers the opportunity to better understand how structural changes propagate from retinal photoreceptors through subcortical relay nuclei to the visual pathways.

In the recent years many therapeutic approaches have been tested preclinically and clinically. However, the majority of the clinical trials fail to restore meaningful vision in patients. Among the multiple factors contributing to the failure of meeting clinical endpoints ^29^, one important consideration is whether, after functional restoration of degenerated or non-functional photoreceptors, the downstream synapses and neurons retain the ability to process and transmit the visual information.

In this study, we adopt a multimodal approach combining high-resolution OCT of retinal layers and fixel-based analysis of white matter tracts to investigate the relationship between retinal remodeling, particularly in the post-receptoral layers, and potential white matter alterations in genetically characterized RP patients. Our goal is to explore whether changes observed in the retina are linked to micro- and macrostructural alterations along the visual pathway, providing insights into potential brain remodeling processes associated with this inherited retinal dystrophy. We focused on PDE6A, PDE6B and RHO-linked RP, as these phenotypes typically represent a retina with relatively well functioning macular cones in the childhood, slow cone degeneration and preservation of foveal structure into adulthood.

## Results

To investigate structural biomarkers along the visual pathway in genetically distinct RP subtypes (RHO, PDE6A, PDE6B), we used a multimodal neuroimaging approach combining high-resolution SD-OCT with diffusion-weighted MRI. Our analysis covered the entire retinocortical pathway, from the segmentation of individual retinal layers to the assessment of white matter structure using fixel-based analysis. A total of 15 RP patients were examined; the foveal region includes data from 14 eyes (instead of 15 for the parafoveal region) due to the presence of a foveal cyst in one patient’s eye, which prevented accurate delineation of the profile.

### Thickness changes of retinal layers

The average of the retinal layers’ thickness at the level of the fovea ad periphery are displayed in Figure 1B-C. Retinal profiles represent the group-averaged values across all participants and genetic subgroups. The shaded areas around the lines indicate the standard error of the mean (SEM).After correcting for the false positive rate (FDR), foveal measurements of RP patients (n = 14) revealed a significant overall loss of retinal thickness of −16.87 ± 14.50% (t = 4.19, P_FDR_ =.0023) compared to controls (t = 4.19, P_FDR_ = 0.0013) (Figure 2A). The greatest foveal loss occurred in the ONL (−55.21 ± 33.34%; t = 5.97, P_FDR_ <.0001) and in the PhL (−32.58 ± 21.45%; t = 5.47, P_FDR_ <.0001). In contrast, several inner retinal layers were significantly thicker than controls, including the RNFL (+34.76 ± 45.90%; t=-2.73, P_FDR_ = 0.017), the IPL (+29.49 ± 27.32%; t = 3.89, P_FDR_ = 0.001) and the INL (+40.30 ± 27.04%; t = −5.37, P_FDR_ <.0001). GCL and OPL didn’t show statistically significant change in comparison to controls.

**Figure 1.**
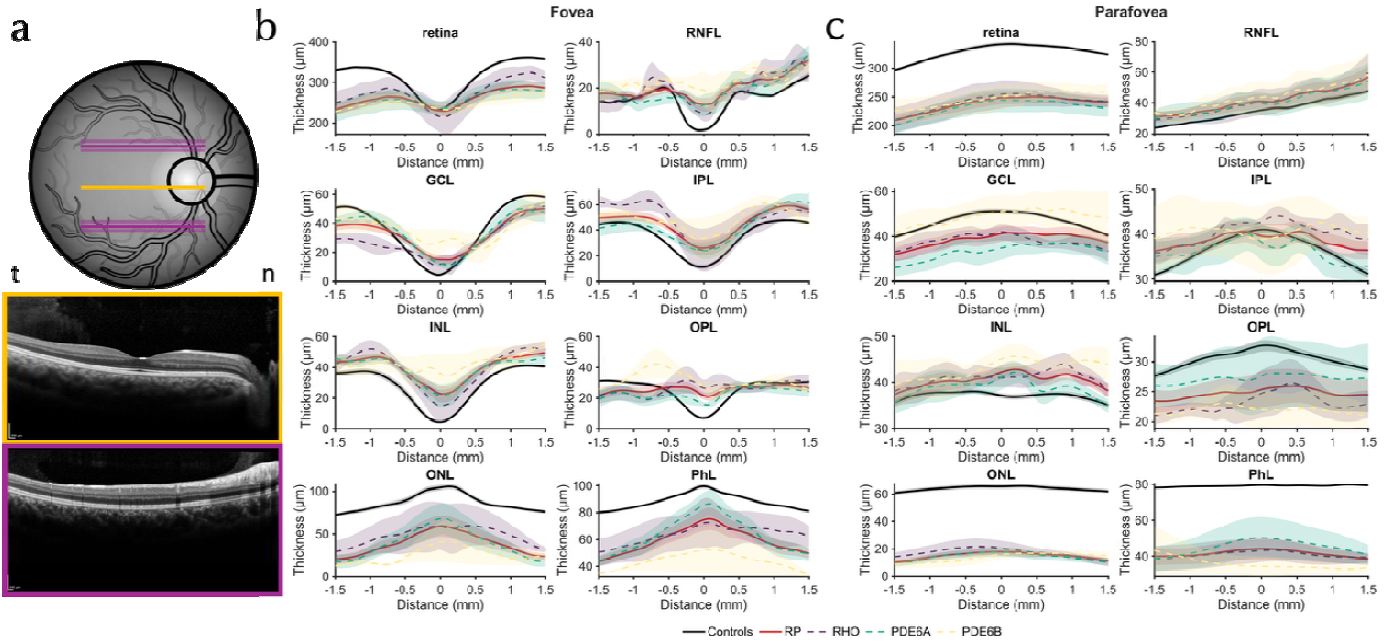
Retinal layer thickness profiles across the fovea and parafovea in control and disease groups. **(a)** Schematic representation of a healthy fundus showing the locations of the three horizontal scanning lines analyzed: one through the fovea (orange line) and one through the parafovea (purple line). Shaded purple parafoveal lines indicate the adjacent scans that were averaged together to have a more precise estimation of the profiles and to account for variability. OCT B-scan images corresponding to each scanning location are displayed below. Temporal and nasal are indicated with t and n, respectively. **(b)** Retinal layer thickness measurements as a function of distance from the foveal center in the foveal region over a 3 mm analyzed area (±1.5 mm from the fovea, representing approximately 6°). Averages of thickness profiles for retinal nerve fiber layer (RNFL), ganglion cell layer (GCL), inner plexiform layer (IPL), inner nuclear layer (INL), outer plexiform layer (OPL), outer nuclear layer (ONL), and photoreceptor layer (PhL) are shown for control subjects (black line) and three disease groups: retinitis pigmentosa (RP, red line), RHO (dashed purple line), PDE6A (dashed cyan line), and PDE6B (dashed yellow line). Shaded regions represent the standard error of the mean (SEM). **(c)** Corresponding retinal layer thickness profiles in the parafoveal region over the same 3 mm analyzed area. All measurements are expressed in micrometers (μm) and plotted relative to distance from the scan center (mm).

**Figure 2.**
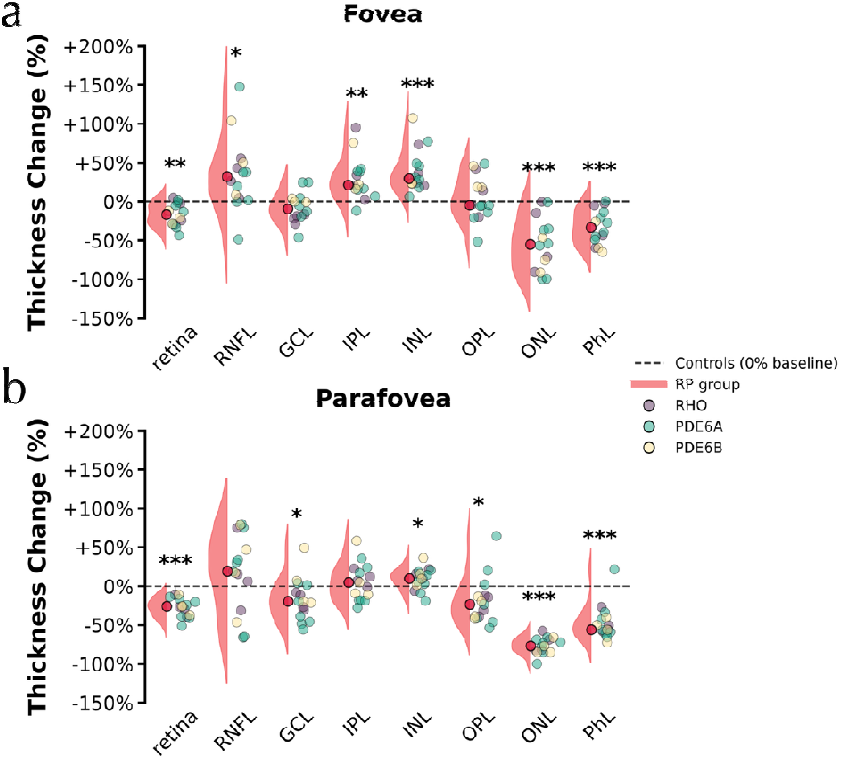
Percentage change in retinal layers thickness in retinitis pigmentosa compared to controls at the fovea and parafovea regions. **a**) Foveal retinal layer thickness changes. **b**) Parafoveal retinal layer thickness changes. Half-violin plots represent the distribution of the entire RP cohort, with the median indicated by a red dot. RP subgroups are stratified by genetic mutation: RHO (purple), PDE6A (green), and PDE6B (yellow). Data are expressed as percentage change relative to healthy controls indicated by the dashed black line. Statistical significance is indicated by asterisks: * P_FDR_ < 0.05; ** P_FDR_ < 0.01; *** P_FDR_ < 0.001. Abbreviations: RNFL, retinal nerve fiber layer; GCL, ganglion cell layer; IPL, inner plexiform layer; INL, inner nuclear layer; OPL, outer plexiform layer; ONL, outer nuclear layer; PhL, photoreceptor layer.

The parafovea retinal area of RP patients (n = 15) exhibited marked thinning, particularly in the total retina (−27.57%; t = 8.815, p <.0001), OPL (–19.23%; t = 2.44; P_FDR_ <.05), ONL (–76.54%; t = 27.79, P_FDR_ <.0001), and PhL (–47.28%; t = 8.12, P_FDR_ <.0001). Moreover, the parafoveal region (Figure 2B) showed a significant decrease of the GCL (–17.67%; t = 2.56; P_FDR_ <.05). An increasing of retinal thickness was found significant just at the level of the INL (+9.19%; t = −2.55, P_FDR_ <.05). RNFL and IPL didn’t show significant changes.

At all regions, ONL thinning was the most consistent and statistically significant finding, accompanied by a significant loss of PhL, while several inner retinal layers (RNFL, IPL, INL) showed thickening especially at the fovea level.

### Decreased fiber density of the white matter tracts

To assess white matter integrity along the visual pathway, we performed fixel-based analysis comparing RP patients and healthy controls across the optic radiation and optic tract defined by tractography (Figure 3A). The analysis was made to find significant changes of white matter properties in RP patients across multiple metrics given the degenerative condition. In Figure 3B, fiber density (FD), was significantly reduced in both the optic radiation (t = 2.981, P_FDR_=.01) and optic tract (t = 5.614, P_FDR_ <.0001), with the optic tract showing stronger diminished FD. The combined measure of fiber density and cross-section (FDC) showed also pronounced differences, with significant reductions observed just at the optic tract level (t = 5.944, P_FDR_ <.0001). Optic radiation, however, showed also a reduction, but didn’t reach the statistical significance (t = 2.191, P_FDR_ = 0.052). Lastly, log fiber-bundle cross-section (log(FC)), didn’t show statistically significant changes, although optic tract (t = 1.795, P_FDR_ = 0.097) resulted to be more reduced than the optic radiation (t = 1.032, P_FDR_ = 0.308) in RP when compared to controls (not shown in the Figure 3). Overall, the analysis of white matter metrics resulted to impact more on the optic tracts compared to the optic radiation, in particular regarding the fiber density parameter.

**Figure 3.**
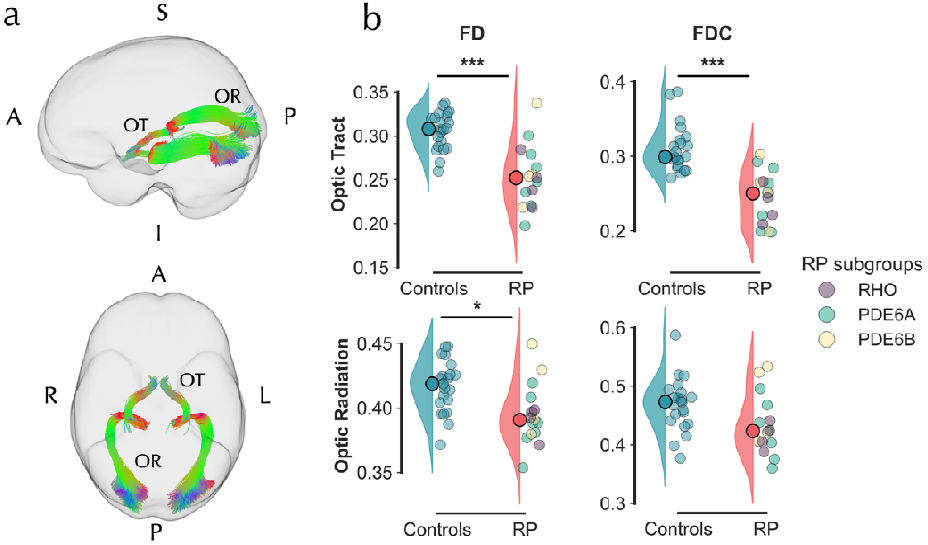
White matter tract reconstruction and fixel metrics analysis in RP. a) Tractography of white matter bundles in MNI template space: optic tracts (OT) and optic radiations (OR) are displayed on sagittal (top) and axial (bottom) views. b) Fixel-Based Analysis (FBA) metrics comparing RP patients and healthy controls. FD, fiber density; FDC, fiber density cross-section. Top row shows OT measurements; bottom row shows OR measurements. RP subgroups are colored with purple (RHO), green (PDE6A) and yellow (PDE6B). Group distributions are shown as half-violin plots, with median values indicated by dots in corresponding saturated colors. Statistical significance after FDR correction is denoted by asterisks (* = P_FDR_<.05; ** = P_FDR_ <.01; *** = P_FDR_ <.001). Abbreviations (panel a): A, anterior; P, posterior; L, left; R, right; OR, optic radiation; OT, optic tract.

### OCT and FBA metrics correlation

Given the observed alterations in both retinal layer thickness and white matter microstructure, we performed a correlation analysis to examine potential relationships between OCT-derived retinal metrics and fixel-based white matter measures. Pearson correlation coefficients were computed to assess the linear association between retinal layer thickness and FBA metrics across the tracts of interest (TOI).

Significant correlations were primarily observed within the IPL layer (Figure 4). In the parafoveal region, IPL thickness showed strong positive correlations with both OR and OT. Specifically, IPL thickness correlated with OR FD (r=0.768, P_FDR_ =.026) whereas FDC resulted not significant (Figure 4B). For the OT, correlations were stronger, with r=0.856 (P_FDR_ =.004) for FD and r=0.801 (P_FDR_=.016) for FDC. Half-violins display group distributions along the respective axes, with IPL thickness on the x-axis and FBA metrics on the y-axis. Notably, in the parafoveal region (Figure 2B), IPL thickness did not differ significantly between patients and controls, whereas FBA metrics showed marked group differences (Figure 3B).

**Figure 4.**
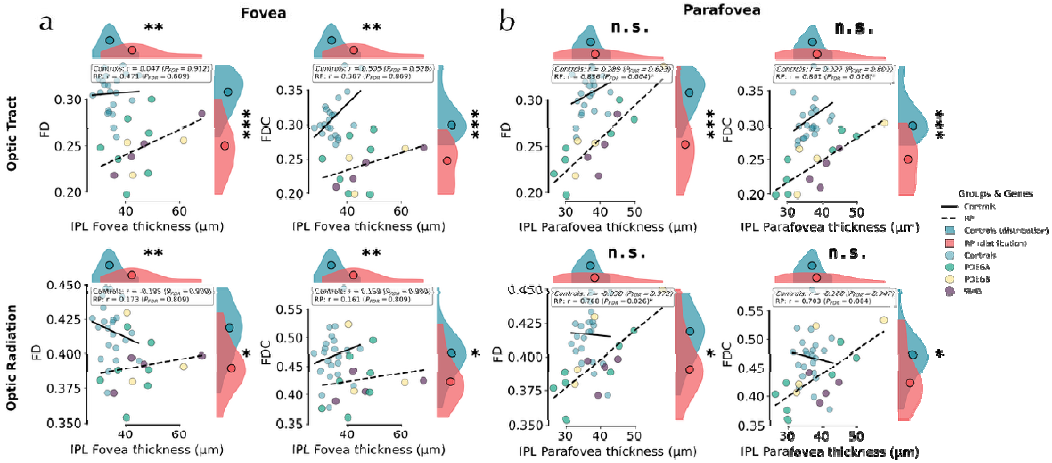
Correlation between IPL retinal layer thickness and white matter fiber metrics in the fovea and parafovea. Scatter plots show the relationship between inner plexiform layer (IPL) thickness (x-axis, µm) and fixel-based analysis (FBA) metrics (y-axis) in RP patients and controls. (**a**) Fovea; (**b**) Parafovea. Top row: Optic tract (OT); bottom row: Optic radiations (OR). In each panel, first column: fiber density (FD); second column: fiber density cross-section (FDC). Controls are shown as light blue circles; RP subtypes as circles: RHO (purple), PDE6A (green), and PDE6B (yellow). Half-violin plots indicate distributions for controls (blue) and RP (red), with median dots. Solid and dashed lines represent linear fits for controls and RP, respectively. Pearson’s r and FDR-corrected p-values are reported; Statistical significance is indicated by asterisks: * P_FDR_ < 0.05; ** P_FDR_ < 0.01; *** P_FDR_ < 0.001.

In the foveal region (Figure 4A), correlations were generally weaker and did not survive correction for multiple comparisons for any TOI. No significant correlations were detected in the control group. In this region, IPL thickness was significantly higher in RP patients compared with controls (Figure 2A), whereas FBA metrics, including log(FC), did not show significant group effects. Notably, in the parafovea (where photoreceptor degeneration is pronounced) IPL thickness values within the range observed in controls were associated with marked reductions in FD, resulting in statistically significant IPL–FD correlations. On the other hand, in the fovea (where photoreceptor layers continue to provide some functional input) an increased thickening of the IPL relative to controls did not translate into a significant association with FD reductions in either the OT or OR. All the correlation results between FBA metrics and retinal layers are reported in the Supplementary Tables 1 and 2.

## Discussion

In this study, we investigated structural alterations along the visual pathway in genetically characterized RP patients by combining high-resolution SD-OCT with fixel-based analysis of diffusion MRI. Our multimodal approach allowed us to assess both retinal layer remodeling and white matter micro/macro structures across the entire retinocortical pathway.

We observed a complex pattern of retinal changes, with marked thinning of the outer layers (ONL, PhL) and relative thickening of several inner layers (RNFL, INL, IPL) in the fovea, whereas the parafoveal retinal area showed more widespread thinning across most layers, with limited preservation of the INL and a significant reduction of the GCL. Within the central visual pathway, fixel-based analysis revealed a predominant reduction of fiber density (FD), particularly in the optic tracts and to a lesser extent in the optic radiations, while macrostructural alterations in fiber-bundle cross-section (log(FC)) were not pronounced. Finally, correlation analyses demonstrated significant associations between retinal and white matter structure, with IPL thickness at parafovea area showing the strongest positive relationships with FD in both the optic tracts and optic radiations. Similar correlations were not observed at the foveal level. Together, these findings highlight distinct regional patterns of retinal remodeling and suggest potential structural links between post-receptoral retinal changes and white matter tracts along the visual pathway.

### Retinal remodeling

Retinal thickness is overall reduced in RP patients. Specifically, our data demonstrate foveal thinning of the photoreceptor layer and thickening of inner post-receptoral layers. The parafoveal region exhibited near-complete absence of outer layers, while inner layers showed a tendency to return toward normal thickness ranges.

The process of post-receptoral thickening has been widely reported ^7,10,12,30-33^, though inconsistent findings for RNFL and GCL suggest variable remodeling patterns ^30,31^. It is known that rod photoreceptor loss initiates retinal remodeling of the inner layers, which, unlike the outer nuclear layer, do not undergo cell death but often remain preserved ^13^. Several mechanisms have been proposed to explain this remodeling. Müller cell dysfunction following photoreceptor degeneration may lead to reactive gliosis and metabolic stress, resulting in neuronal swelling and thickening of the inner retina ^34^. This hypothesis is supported by parallels with other conditions: for instance, inner retinal thickening is also observed in Behçet’s inflammatory disease ^35^, while an increase in aqueous flare associated with photoreceptor loss suggests a link to intraocular inflammation ^33^. In addition to metabolic stress and inflammation, mechanical factors have also been suggested. Axonal swelling, structural remodeling involving neuronal migration, and glial hypertrophy may all contribute to increased thickness ^7,36^. Although the present study was not designed to identify the primary cause of post-receptoral thickening, our observations provide some insights. In our data, the pattern of inner retinal changes argues against a purely mechanical explanation. The parafoveal region, where outer layers were largely absent, did not exhibit a corresponding thickening of the inner retina. In the foveal region, the thickening was most pronounced where the photoreceptor layer was still partially preserved. This could indicate that post-receptoral thickening is not merely a geometric or compensatory response to outer retinal loss, but rather reflects active cellular remodeling, potentially related to Müller cell gliosis or metabolic alterations ^37^.

### Fibre density reduction in optic tracts and radiations

Our results show that the optic tracts and optic radiations exhibited a marked reduction in fibre density (FD), indicating axonal loss without accompanying reductions in fibre bundle size or atrophy. The additional significant difference observed in fibre density and cross-section (FDC) appears to be primarily driven by the reduction in FD alone. In the context of FBA and eye diseases, Haykal et al. (2019) showed a reduction of both FD and FC in the optic tracts, while in the optic radiations the effect was confined to FD ^27^. This pattern suggests a temporal sequence, in which axonal density loss represents an early event, followed by fiber atrophy in structures more proximal to the optic nerve. A similar phenomenon has been described in optic neuritis associated with multiple sclerosis (MS). Using FBA, Gajamange et al. (2018) reported a relatively specific reduction of FD, and to a lesser extent FC in the visual-ralated tracts ^25^. Accordingly, they concluded that an initial FD reduction represents the earliest manifestation of damage, followed later by a decline in FC, with FDC integrating both the micro- and macrostructural dimensions of degeneration.

In the context of our results, the preferential FD loss in the optic tracts and radiations likely reflects an early disruption of visual system integrity. At later disease stages, progressive deafferentation may also lead to marked tract atrophy. It remains to be determined whether FD reduction represents a reversible functional disconnection (potentially recoverable with restored retinal input) or a structural loss of axonal efficiency with limited recovery potential.

### Retina-visual tracts relationship

Previous studies investigating white matter tracts in RP populations remain inconclusive as to whether retinal degeneration systematically leads to downstream alterations within the central visual system ^20,38,39^. Anterograde trans-synaptic degeneration describes progressive downstream involvement of central visual pathways following retinal pathology, while retrograde degeneration refers to upstream neuronal loss. In the anterograde model, loss of retinal ganglion cells (RGCs) and their axons projecting to the LGN may sequentially lead to optic tract and radiation alterations. Such associations between RGC thinning and white matter changes have been reported in Leber’s hereditary optic neuropathy (LHON) ^40-42^, glaucoma ^43^ and optic neuritis ^44^. In photoreceptor-driven dystrophies, Ogawa and colleagues ^40^ reported alterations in diffusion parameters within the optic tracts and radiations, in the absence of changes in RNFL thickness. The authors suggested that central white matter degeneration in cone-rod dystrophy (CRD) patients may follow mechanisms that are partially independent from the retinal alterations. Along the same line, in age-related macular degeneration (AMD), a previous study demonstrated an association between reduced RNFL thickness and alterations in the optic tracts, whereas no such relationship was detected with the optic radiations ^45^. These observations support the view that different portions of the visual pathway may be differentially affected, and that central white matter involvement does not always directly mirror retinal structural loss.

In our cohort, we observed thickness differences across foveal and parafoveal retinal layers. A notable finding was that parafoveal inner retinal thickening, while still greater than controls, approached normal levels. This pattern suggests an adaptive remodeling response, where residual photoreceptors and post-receptoral layers maintain activity and functional connections with central visual tracts. In the parafovea, however, although the IPL remained significantly thicker than controls, the overall inner retinal thickness showed a trend toward normalization. Importantly, this partial return to normal thickness correlated with reduced fibre density in optic tracts and radiations. We interpret this as a degeneration post-receptoral process, in which the initial hypertrophic remodeling of the inner retina loses its adaptive role, coinciding with measurable downstream axonal loss.

### Limitations

Several methodological considerations should be acknowledged. First, the patient and control groups were not fully matched for age, which may have introduced residual confounding effects on diffusion-derived measures and retinal layer thickness, both known to vary with aging. Second, OCT data were acquired from a single eye per patient, limiting the ability to assess variability and potential asymmetries in retinal degeneration and its central correlates. Nonetheless, patients with RP generally exhibit a high degree of symmetry between eyes ^46-48^, suggesting that measurements from a single eye can still provide a representative estimate of the disease severity. Bilateral imaging would allow for a more comprehensive characterization of structural relationships across the visual pathway. Additionally, the sample size was modest, particularly given the rarity and genetic heterogeneity of RP, which may have reduced sensitivity to subtle effects and limited generalizability. Finally, the cross-sectional design prevents firm conclusions about the temporal sequence of retinal and white matter changes; longitudinal assessments are needed to determine whether white matter alterations evolve secondarily to retinal degeneration or reflect parallel disease processes.

## Conclusions

In this study, we investigated three genetic subtypes of retinitis pigmentosa (RHO, PDE6A, and PDE6B; total = 15 eyes) and healthy controls (total = 22 eyes) using a multimodal approach combining retinal morphological characterization (OCT) with diffusion-weighted imaging (DWI) of the visual pathway white matter tracts. Our goal was to identify potential links between the photoreceptor degeneration occurring in the retina and its possible propagation along the visual pathway.

Our findings show that the near-complete loss of the photoreceptor layer, predominantly in the parafoveal region, is not accompanied by a linear degeneration of the post-receptoral retinal layers. On the contrary, we observed a thickening of the inner retinal layers in the foveal region, where photoreceptors are relatively preserved, and near-normal thickness in regions where photoreceptors are absent. This phenomenon likely reflects retinal remodeling secondary to the disease, which we attribute to Müller cell gliosis rather than to a mechanical compensatory mechanism driven by remaining cells occupying the degenerated space. In parallel, fixel-based analysis of the DWI data revealed that this retinal remodeling is associated with microstructural alterations in the visual pathway, characterized by reduced fiber density without corresponding macrostructural changes in tract volume. The effect was more pronounced in the optic tracts than in the optic radiations, supporting the hypothesis of trans-synaptic deafferentation along the visual pathway.

It remains unclear whether the reduced fiber density still allows for normal functional signal transmission in the context of retinal interventions. Moreover, whether therapeutic treatments aimed at restoring photoreceptor function could also promote re-afferentation of these microstructural pathways remains unknown. Applying this multimodal framework in a post-intervention context will be crucial to evaluate potential recovery mechanisms along the visual pathway.

## Materials and Methods

### Participants

A total of 37 participants took part of the study, comprising 22 healthy controls (mean age = 25.36 ± 4.9 years; 12 females, 10 males) and 15 patients with a confirmed genetic diagnosis of retinitis pigmentosa (RP) (mean age = 36.9 ± 11.5 years; 4 females, 11 males). Retinal data were collected from one eye per RP participant (right eye; *n* = 15). RP patients were recruited through the University Eye Hospital in Tübingen based on their mutation. The following genotypes were included for the study: 4 patients with RHO, 6 with PDE6A, and 5 with PDE6B. Healthy control participants were selected based on the absence of any retinal pathology and had normal visual acuity. The mean (SD) BCVA across 15 eyes with RP was 0.26 (0.30) logMAR (range, −0.10 to 1.10; Snellen equivalent ≈ 20/36; 0.65 decimal), indicating mild-to-moderate visual acuity reduction. One patient presented with a foveal cyst; data from this eye were excluded from the foveal analyses but kept for parafoveal analyses. All participants were volunteers and provided written informed consent prior to participation, in accordance with the Declaration of Helsinki. The study was approved by the Ethics Board of the University of Tübingen under the number 884/2020BO2.

### Retinal images acquisition

Volumetric retinal imaging was performed using spectral-domain optical coherence tomography (SD-OCT) with the Spectralis HDR + OCT instrument (Heidelberg Engineering GmbH, Heidelberg, Germany). Each volumetric scan consisted of 31 B-scans, with 768 A-scans per B-scan and 496 axial samples per A-scan, centered on the macula. The axial resolution was 3.9 µm. The acquisition included automatic real-time (ART) averaging, with the number of averages per B-scan ranging from 28 to 31. All scans were acquired in high-resolution mode using the macular fixation target and were centered on the fovea. For each subject, retinal layers were manually segmented using the OCT software (HEYEX v. 2.5.9; Heidelberg Engineering) and the resulting data were exported and imported into MATLAB (R2023b, The MathWorks, Natick, Massachusetts). The layers extracted included: the whole retina (from internal limiting membrane to Bruch’s membrane), ganglion cell layer (GCL), inner plexiform layer (IPL), inner nuclear layer (INL), outer plexiform layer (OPL) and outer nuclear layer (ONL). Three retinal regions were analyzed within a 6° area around the fovea (corresponding to a 3 mm diameter): the fovea and the superior and inferior parafoveal regions (each 3 mm above and below the foveal center). For each parafoveal region, the mean of three adjacent B-scans was computed to obtain representative layer thickness values.

### Brain images acquisition

#### Structural images

High-resolution T1-weighted and T2-weighted structural images were acquired using a 3T Siemens Prisma scanner (Siemens Healthcare, Erlangen, Germany) equipped with a 64-channel Head/Neck coil at the University Eye Hospital in Tübingen, Germany. The T1-weighted images were obtained using a 3D magnetization-prepared rapid gradient echo (MPRAGE) sequence with the following parameters: repetition time (TR) = 2400 ms, echo time (TE) = 2.22 ms, inversion time (TI) = 1000 ms, flip angle = 8°, slice thickness = 0.8 mm, parallel imaging (GRAPPA) with an acceleration factor of 2, and a matrix size of 320 × 300 with a base resolution of 320. The T2-weighted images were acquired using a 3D sampling perfection with application-optimized contrasts using different flip angle evolution (SPACE) sequence with TR = 3200 ms, TE = 563 ms, flip angle = 120°, slice thickness = 0.8 mm, GRAPPA acceleration factor of 2, and matrix size of 320 × 300. Both sequences were acquired with full k-space coverage and 100% sampling, and images were oriented sagittally with partial Fourier sampling and no nonlinear gradient correction. The acquired data were later converted to NIfTI format using *dcm2niix* (version v1.0.20220720).

#### Diffusion weighted images

Diffusion weighted imaging (DWI) data were acquired on a 3T Siemens Prisma MRI scanner (Siemens Healthcare, Erlangen, Germany) using syngo MR E11 software. Diffusion-weighted images were obtained using a multiband echo-planar imaging sequence with 99 diffusion gradient directions in the posterior-anterior (PA) direction and corresponding images in the anterior-posterior (AP) direction. Two DWI shells were acquired, b=1500 s/mm^2^, and b=3000 s/mm^2^ and seven images with no diffusion weighting (b=0 s/mm2) were also acquired in each phase encoding direction. Other acquisition parameters included: TR = 3230 ms, TE = 89 ms, flip angle = 78° and voxel size of 1.5 mm^3^. Single-band reference images were also obtained for each phase encoding direction to facilitate subsequent motion and distortion correction procedures. Structural and DWI data were organized according to the Brain Imaging Data Structure (BIDS) format.

### Preprocessing of Imaging

#### Diffusion data preprocessing

Preprocessing was performed using *QSIPrep* 0.20.1 ^49^, which is based on Nipype 1.8.6 ^50^. Many internal operations of QSIPrep use Nilearn 0.10.2 ^51^ and Dipy ^52^.

#### Anatomical images

The T1-weighted (T1w) image was corrected for intensity non-uniformity (INU) using N4BiasFieldCorrection (ANTs 2.4.3) ^53^ and used as an anatomical reference throughout the workflow. The anatomical reference image was reoriented into AC-PC alignment via a 6-DOF transform extracted from a full Affine registration to the MNI152NLin2009cAsym template. A full nonlinear registration to the template from AC-PC space was estimated via symmetric nonlinear registration (SyN) using antsRegistration. Brain extraction was performed on the T1w image using SynthStrip ^54^ and automated segmentation was performed using SynthSeg (^55^, @synthseg2) from Freesurfer version 7.3.1. Additionally, structural images were segmented using Freesurfer *recon-all* command. The Glasser HCP-MMP1 atlas ^56^ was then applied to each subject by projecting the *fsaverage* annotation files to the individual cortical surfaces using *mri_surf2surf* for both hemispheres. Parcel-wise cortical morphometry (thickness, surface area, and volume) was then extracted with *mris_anatomical_stats*. The surface parcellation was converted into a volumetric segmentation in the subject’s native space using *mri_aparc2aseg* and *labelconvert* to ensure a consistent labeling scheme across subjects. The resulting parcellations were available in both surface-based and volumetric formats for subsequent analyses. Finally, the lateral geniculate nucleus (LGN) was extracted by segmenting the thalamus into its constituent nuclei using the *segmentThalamicNuclei*.*sh* script from Freesurfer. This method performs a Bayesian segmentation into 25 nuclei based on histological probability atlases ^57^.

#### Diffusion images

Diffusion images were grouped into two phase encoding polarity groups. Any images with a b-value less than 100 s/mm^2 were treated as a b=0 image. MP-PCA denoising as implemented in MRtrix3’s *dwidenoise* ^58^ was applied with a 5-voxel window. After MP-PCA, Gibbs unringing was performed using MRtrix3’s *mrdegibbs* ^59^. Following unringing, the mean intensity of the DWI series was adjusted so all the mean intensity of the b=0 images matched across each separate DWI scanning sequence. B1 field inhomogeneity was corrected using *dwibiascorrect* from MRtrix3 with the N4 algorithm ^53^ after corrected images were resampled. Both distortion groups were then merged into a single file, as required for the FSL workflows.

FSL (version 6.0.5.1:57b01774)’s *eddy* was used for head motion correction and Eddy current correction ^60^. Eddy was configured with a q-space smoothing factor of 10, a total of 5 iterations, and 1000 voxels used to estimate hyperparameters. A linear first level model and a linear second level model were used to characterize Eddy current-related spatial distortion. q-space coordinates were forcefully assigned to shells. Field offset was attempted to be separated from subject movement. Shells were aligned post-eddy. Eddy’s outlier replacement was run ^61^. Data were grouped by slice, only including values from slices determined to contain at least 250 intracerebral voxels. Groups deviating by more than 4 standard deviations from the prediction had their data replaced with imputed values.

Data was collected with reversed phase-encode blips, resulting in pairs of images with distortions going in opposite directions. FSL’s TOPUP ^62^ was used to estimate a susceptibility-induced off-resonance field based on b=0 images extracted from multiple DWI series with reversed phase encoding directions. The TOPUP-estimated fieldmap was incorporated into the Eddy current and head motion correction interpolation. Final interpolation was performed using the *jac* method.

Several confounding time-series were calculated based on the preprocessed DWI: framewise displacement using the implementation in Nipype (following the definitions by ^63^). The head-motion estimates calculated in the correction step were also placed within the corresponding confounds file. Slicewise cross correlation was also calculated. The DWI time-series were resampled to AC-PC, generating a preprocessed DWI run in ACPC space with 1.3mm isotropic voxels.

#### Fixel-based analysis

Fixel-based analysis (FBA) was performed using MRtrix3 (www.mrtrix.org) to investigate white matter microstructure and macrostructure. Following preprocessing with *QSIPrep*, DWI data were reoriented to standard MNI space using FSL and upsampled to 1.25 mm isotropic voxels. Multi-shell multi-tissue constrained spherical deconvolution (CSD) was performed using the unsupervised Dhollander algorithm to estimate tissue response functions for white matter, gray matter, and cerebrospinal fluid ^64^. Fiber orientation distributions (FODs) were computed separately for patient and control groups using group-averaged response functions, followed by intensity normalization across tissue compartments.

An unbiased population-specific FOD template was generated from all participants using an iterative registration approach. Individual FODs were then registered to the template using non-linear registration, and fixel metrics were computed in template space. Three complementary fixel metrics were calculated: fiber density (FD), reflecting intra-axonal volume ^65^; fiber-bundle cross-section (FC), indicating macrostructural changes ^22^; and fiber density and cross-section (FDC), representing the product of the two metrics ^22^. We applied a log transform to FC so it would be normally distributed and centered around zero. FDC was calculated before this log transformation was applied.

A whole-brain tractogram of 20 million streamlines was generated from the template FOD and subsequently filtered to 2 million streamlines using spherical-deconvolution informed filtering of tractograms (SIFT) ^66^. Fixel-fixel connectivity matrices were computed to enable smoothing of fixel data along fiber bundles. This connectivity data was used to inform spatial smoothing of FD, log(FC), and FDC maps, such that smoothing at a given fixel only occurred within that fixel’s fiber population, thus avoiding partial-volume effects or influences from crossing fibers ^67^.

#### Tract segmentation

White matter of the optic radiations (OR) was automatically delineated using TractSeg version 2.3 ^68^, a deep learning based method to segment white matter and reconstruct fiber bundles. Conversely, optic tracts (OT) were extracted by seeding two regions of interest based on the template at the level of the optic chiasm and LGN; subsequently, the 2 million streamlines tractography was edited according to the seeded region of interests using *tckedit* command. Tract-specific fixel metrics were then extracted by averaging values within each tract’s fixel mask created for both tracts of interest (TOI), OR and OT. From each set of fiber bundle streamlines, we created a corresponding fixel tract density map, which we binarized to create tract fixel masks.

### Statistical analyses

#### Percentage thickness loss

To quantify retinal degeneration in RP patients, we calculated the percentage thickness loss for each retinal layer (whole retina, RNFL, GCL, PhL, INL, IPL, ONL, OPL) by comparing individual patient values to the average thickness of age-matched healthy controls. This was done across three anatomical regions: the fovea, and the superior and inferior retina at 6 mm eccentricity. Thickness loss was expressed as a percentage using the formula:

*[(mean control thickness – RP patient thickness) /mean control thickness] × 100*.

To determine whether the observed thickness loss in RP patients was statistically significant (i.e., different from 0% loss, which would indicate no difference from controls), we performed one-sample statistical tests for each layer and region. Recognizing that clinical data can deviate from normal distributions, we applied both parametric and non-parametric methods: one-sample t-tests to assess group-level differences and Wilcoxon signed-rank tests as distribution-free alternatives.

Before applying the t-tests, we tested for normality using the Shapiro-Wilk test (α = 0.05) to decide whether a parametric or non-parametric approach was more appropriate in each case.

To correct for multiple comparisons across the various retinal layers and regions, we applied the Benjamini-Hochberg false discovery rate (FDR) correction (α = 0.05), calculating adjusted p-values separately for the performed analyses.

#### FBA metrics

To examine associations between RP patients and controls in both FBA-derived metrics we used both parametric and non-parametric testing. For each tract–metric combination, group differences were assessed with both independent-samples t-tests (parametric) and Mann–Whitney U tests (non-parametric) to account for potential deviations from normality in the data distribution. To correct for multiple comparisons, p-values (α = 0.05) were adjusted using the Benjamini–Hochberg FDR correction applied across all comparisons.

#### OCT – FBA metrics correlation

To explore the relationship between retinal layer thickness and white matter tract structure, we conducted a comprehensive correlation analysis across all combinations of retinal layers, eccentricity regions, FBA diffusion metrics related to OT and OR. For each subject, tract-specific diffusion values were extracted contralateral to the examined eye. Analyses were performed separately for RP patients and healthy controls. For each group, we computed standard Pearson correlations to assess linear relationships. Only combinations with at least three valid data points per group were included. For each correlation, we recorded the number of subjects, mean and standard deviation of age, and the resulting correlation coefficients and p-values. To correct for multiple comparisons across all combinations, we applied the Benjamini-Hochberg FDR correction (α = 0.05) independently to each correlation method. All computations were performed in Python using Scipy and Statsmodels packages.

## Acknowledgements

This study was supported by the Tistou & Charlotte Kerstan foundation project RI-FG P5 1-1.

## Author contributions

RG: Conceptualization, Software, Formal analysis, Visualization, Writing original draft, Writing – Review & Editing

VCV: Investigation, Data curation, Software, Writing – Review & Editing

MR: Investigation (patient recruitment), Writing – Review & Editing

LW: Investigation (patient recruitment), Writing – Review & Editing

LK: Investigation (patient recruitment), Writing – Review & Editing

DM: Investigation (patient recruitment), Writing – Review & Editing

CB: Methodology, Supervision, Writing – Review & Editing

KaS: Conceptualization, Supervision, Resources, Project administration, Funding acquisition, Writing original draft, Writing – Review & Editing

KrS: Conceptualization, Supervision, Resources, Writing original draft, Writing – Review & Editing

## Supplementary Materials

### Supplementary Table 1

**Supplemental Table 1.**
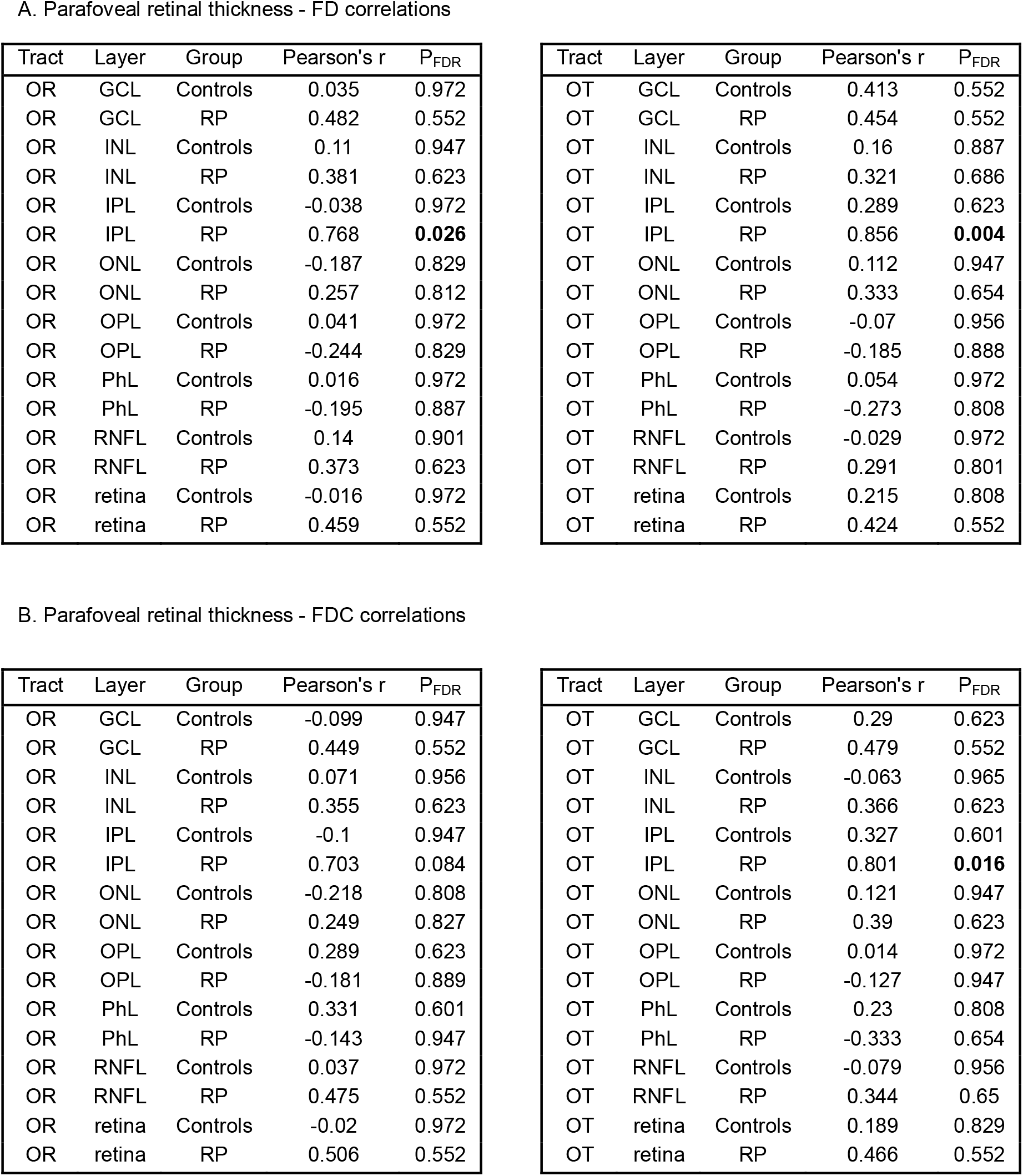
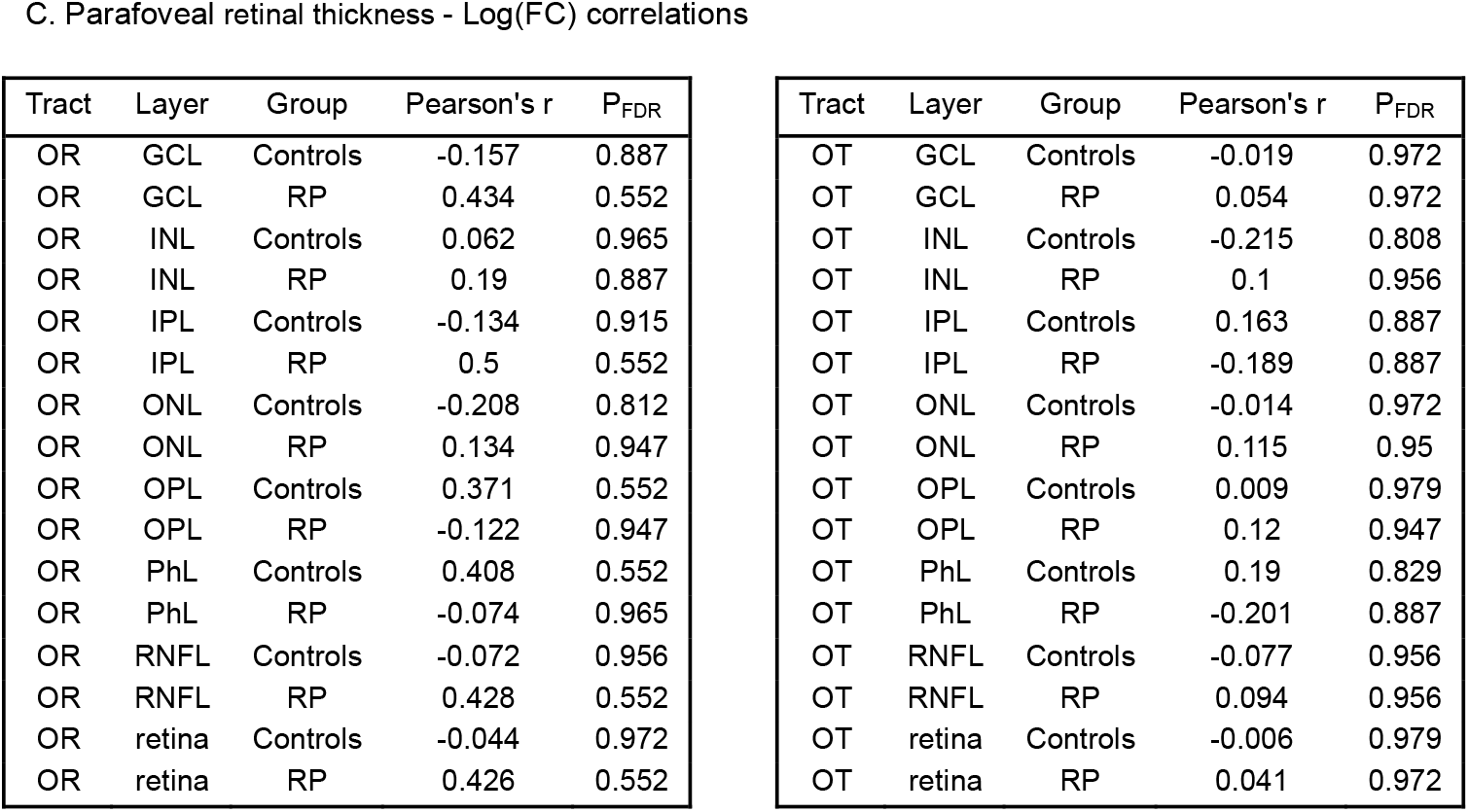
Correlations between parafoveal retinal layer thickness and white matter fibre metrics along the visual pathway in RP patients and controls. The table reports Pearson’s correlation coefficients (*r*) and FDR-corrected *p*-values (*P*_FDR_) for each retinal layer and tract combination. Analyses were performed separately for the optic radiation (OR) and optic tract (OT) in relation to the thickness of individual retinal layers (retinal nerve fibre layer (RNFL), ganglion cell layer (GCL), inner plexiform layer (IPL), inner nuclear layer (INL), outer plexiform layer (OPL), outer nuclear layer (ONL), and photoreceptor layer (PhL)) as well as total retinal thickness. Correlations were computed for both controls and RP patients across three fixel-based metrics: A) fibre density (FD); B) combined fibre density and cross-section (FDC); C) log-transformed fibre cross-section (log(FC)). Significant correlations after FDR correction are indicated in bold (*P*_*FDR*_ < 0.05).

A. Parafoveal retinal thickness - FD correlations
| Tract | Layer | Group | Pearson's r | P<sub>FDR</sub> |
| --- | --- | --- | --- | --- |
| OR | GCL | Controls | 0.035 | 0.972 |
| OR | GCL | RP | 0.482 | 0.552 |
| OR | INL | Controls | 0.11 | 0.947 |
| OR | INL | RP | 0.381 | 0.623 |
| OR | IPL | Controls | -0.038 | 0.972 |
| OR | IPL | RP | 0.768 | **0.026** |
| OR | ONL | Controls | -0.187 | 0.829 |
| OR | ONL | RP | 0.257 | 0.812 |
| OR | OPL | Controls | 0.041 | 0.972 |
| OR | OPL | RP | -0.244 | 0.829 |
| OR | PhL | Controls | 0.016 | 0.972 |
| OR | PhL | RP | -0.195 | 0.887 |
| OR | RNFL | Controls | 0.14 | 0.901 |
| OR | RNFL | RP | 0.373 | 0.623 |
| OR | retina | Controls | -0.016 | 0.972 |
| OR | retina | RP | 0.459 | 0.552 |
| OT | GCL | Controls | 0.413 | 0.552 |
| OT | GCL | RP | 0.454 | 0.552 |
| OT | INL | Controls | 0.16 | 0.887 |
| OT | INL | RP | 0.321 | 0.686 |
| OT | IPL | Controls | 0.289 | 0.623 |
| OT | IPL | RP | 0.856 | **0.004** |
| OT | ONL | Controls | 0.112 | 0.947 |
| OT | ONL | RP | 0.333 | 0.654 |
| OT | OPL | Controls | -0.07 | 0.956 |
| OT | OPL | RP | -0.185 | 0.888 |
| OT | PhL | Controls | 0.054 | 0.972 |
| OT | PhL | RP | -0.273 | 0.808 |
| OT | RNFL | Controls | -0.029 | 0.972 |
| OT | RNFL | RP | 0.291 | 0.801 |
| OT | retina | Controls | 0.215 | 0.808 |
| OT | retina | RP | 0.424 | 0.552 |

B. Parafoveal retinal thickness - FDC correlations
| Tract | Layer | Group | Pearson's r | P<sub>FDR</sub> |
| --- | --- | --- | --- | --- |
| OR | GCL | Controls | -0.099 | 0.947 |
| OR | GCL | RP | 0.449 | 0.552 |
| OR | INL | Controls | 0.071 | 0.956 |
| OR | INL | RP | 0.355 | 0.623 |
| OR | IPL | Controls | -0.1 | 0.947 |
| OR | IPL | RP | 0.703 | 0.084 |
| OR | ONL | Controls | -0.218 | 0.808 |
| OR | ONL | RP | 0.249 | 0.827 |
| OR | OPL | Controls | 0.289 | 0.623 |
| OR | OPL | RP | -0.181 | 0.889 |
| OR | PhL | Controls | 0.331 | 0.601 |
| OR | PhL | RP | -0.143 | 0.947 |
| OR | RNFL | Controls | 0.037 | 0.972 |
| OR | RNFL | RP | 0.475 | 0.552 |
| OR | retina | Controls | -0.02 | 0.972 |
| OR | retina | RP | 0.506 | 0.552 |
| OT | GCL | Controls | 0.29 | 0.623 |
| OT | GCL | RP | 0.479 | 0.552 |
| OT | INL | Controls | -0.063 | 0.965 |
| OT | INL | RP | 0.366 | 0.623 |
| OT | IPL | Controls | 0.327 | 0.601 |
| OT | IPL | RP | 0.801 | **0.016** |
| OT | ONL | Controls | 0.121 | 0.947 |
| OT | ONL | RP | 0.39 | 0.623 |
| OT | OPL | Controls | 0.014 | 0.972 |
| OT | OPL | RP | -0.127 | 0.947 |
| OT | PhL | Controls | 0.23 | 0.808 |
| OT | PhL | RP | -0.333 | 0.654 |
| OT | RNFL | Controls | -0.079 | 0.956 |
| OT | RNFL | RP | 0.344 | 0.65 |
| OT | retina | Controls | 0.189 | 0.829 |
| OT | retina | RP | 0.466 | 0.552 |

Supplemental Table 1. **Correlations between parafoveal retinal layer thickness and white matter fibre metrics along the visual pathway in RP patients and controls.**
| Tract | Layer | Group | Pearson's r | P<sub>FDR</sub> |
| --- | --- | --- | --- | --- |
| OR | GCL | Controls | -0.157 | 0.887 |
| OR | GCL | RP | 0.434 | 0.552 |
| OR | INL | Controls | 0.062 | 0.965 |
| OR | INL | RP | 0.19 | 0.887 |
| OR | IPL | Controls | -0.134 | 0.915 |
| OR | IPL | RP | 0.5 | 0.552 |
| OR | ONL | Controls | -0.208 | 0.812 |
| OR | ONL | RP | 0.134 | 0.947 |
| OR | OPL | Controls | 0.371 | 0.552 |
| OR | OPL | RP | -0.122 | 0.947 |
| OR | PhL | Controls | 0.408 | 0.552 |
| OR | PhL | RP | -0.074 | 0.965 |
| OR | RNFL | Controls | -0.072 | 0.956 |
| OR | RNFL | RP | 0.428 | 0.552 |
| OR | retina | Controls | -0.044 | 0.972 |
| OR | retina | RP | 0.426 | 0.552 |
| OT | GCL | Controls | -0.019 | 0.972 |
| OT | GCL | RP | 0.054 | 0.972 |
| OT | INL | Controls | -0.215 | 0.808 |
| OT | INL | RP | 0.1 | 0.956 |
| OT | IPL | Controls | 0.163 | 0.887 |
| OT | IPL | RP | -0.189 | 0.887 |
| OT | ONL | Controls | -0.014 | 0.972 |
| OT | ONL | RP | 0.115 | 0.95 |
| OT | OPL | Controls | 0.009 | 0.979 |
| OT | OPL | RP | 0.12 | 0.947 |
| OT | PhL | Controls | 0.19 | 0.829 |
| OT | PhL | RP | -0.201 | 0.887 |
| OT | RNFL | Controls | -0.077 | 0.956 |
| OT | RNFL | RP | 0.094 | 0.956 |
| OT | retina | Controls | -0.006 | 0.979 |
| OT | retina | RP | 0.041 | 0.972 |

### Supplementary Table 2

**Supplemental Table 2.**
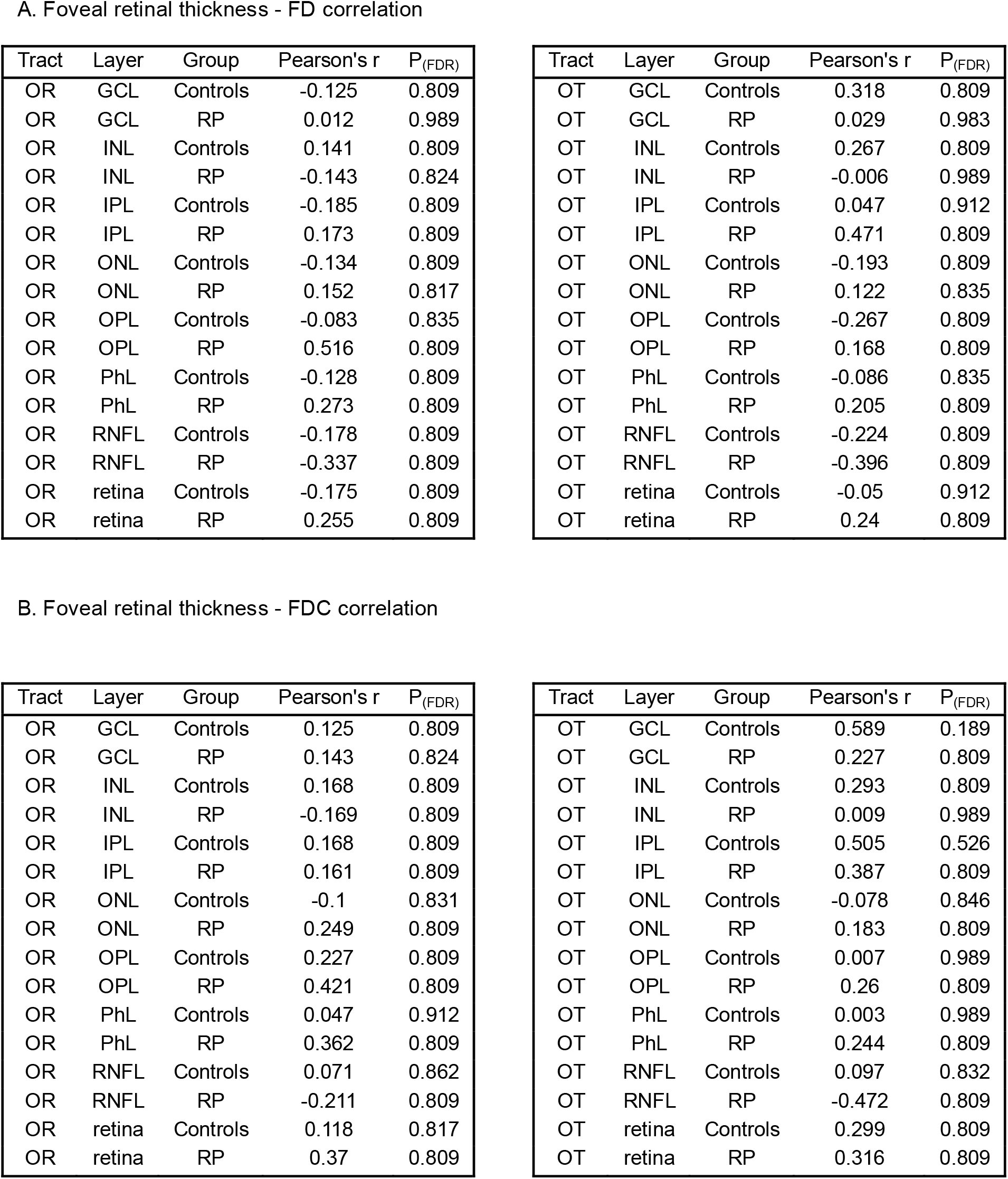
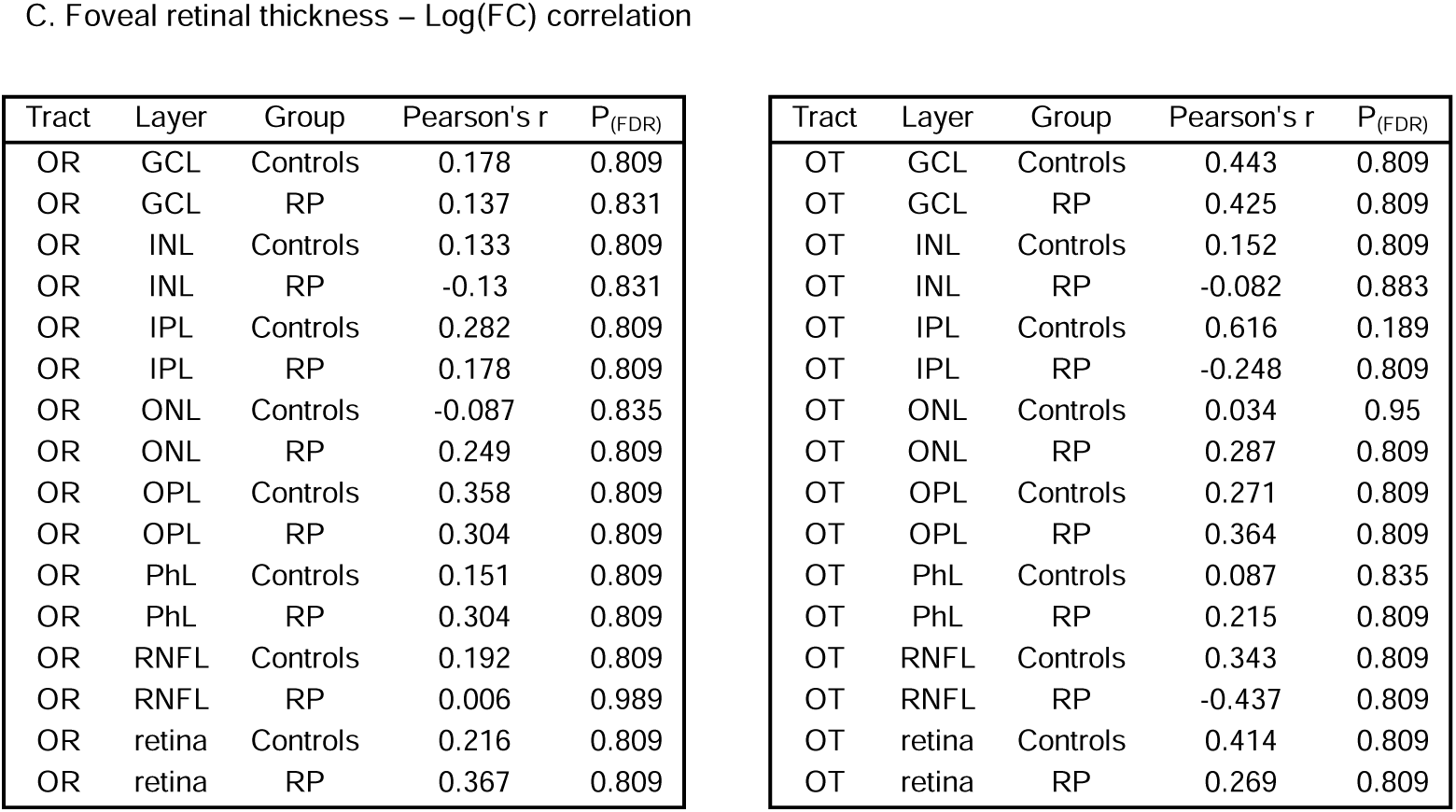
Correlations between foveal retinal layer thickness and white matter fibre metrics along the visual pathway in RP patients and controls. The table reports Pearson’s correlation coefficients (*r*) and FDR-corrected *p*-values (*P*_FDR_) for each retinal layer and tract combination. Analyses were performed separately for the optic radiation (OR) and optic tract (OT) in relation to the thickness of individual retinal layers (retinal nerve fibre layer (RNFL), ganglion cell layer (GCL), inner plexiform layer (IPL), inner nuclear layer (INL), outer plexiform layer (OPL), outer nuclear layer (ONL), and photoreceptor layer (PhL)) as well as total retinal thickness. Correlations were computed for both controls and RP patients across three fixel-based metrics: A) fibre density (FD); B) combined fibre density and cross-section (FDC); C) log-transformed fibre cross-section (log(FC)). No correlations were found significant.

A. Foveal retinal thickness - FD correlation
| Tract | Layer | Group | Pearson's r | P<sub>(FDR)</sub> |
| --- | --- | --- | --- | --- |
| OR | GCL | Controls | -0.125 | 0.809 |
| OR | GCL | RP | 0.012 | 0.989 |
| OR | INL | Controls | 0.141 | 0.809 |
| OR | INL | RP | -0.143 | 0.824 |
| OR | IPL | Controls | -0.185 | 0.809 |
| OR | IPL | RP | 0.173 | 0.809 |
| OR | ONL | Controls | -0.134 | 0.809 |
| OR | ONL | RP | 0.152 | 0.817 |
| OR | OPL | Controls | -0.083 | 0.835 |
| OR | OPL | RP | 0.516 | 0.809 |
| OR | PhL | Controls | -0.128 | 0.809 |
| OR | PhL | RP | 0.273 | 0.809 |
| OR | RNFL | Controls | -0.178 | 0.809 |
| OR | RNFL | RP | -0.337 | 0.809 |
| OR | retina | Controls | -0.175 | 0.809 |
| OR | retina | RP | 0.255 | 0.809 |
| OT | GCL | Controls | 0.318 | 0.809 |
| OT | GCL | RP | 0.029 | 0.983 |
| OT | INL | Controls | 0.267 | 0.809 |
| OT | INL | RP | -0.006 | 0.989 |
| OT | IPL | Controls | 0.047 | 0.912 |
| OT | IPL | RP | 0.471 | 0.809 |
| OT | ONL | Controls | -0.193 | 0.809 |
| OT | ONL | RP | 0.122 | 0.835 |
| OT | OPL | Controls | -0.267 | 0.809 |
| OT | OPL | RP | 0.168 | 0.809 |
| OT | PhL | Controls | -0.086 | 0.835 |
| OT | PhL | RP | 0.205 | 0.809 |
| OT | RNFL | Controls | -0.224 | 0.809 |
| OT | RNFL | RP | -0.396 | 0.809 |
| OT | retina | Controls | -0.05 | 0.912 |
| OT | retina | RP | 0.24 | 0.809 |

B. Foveal retinal thickness - FDC correlation
| Tract | Layer | Group | Pearson's r | P<sub>(FDR)</sub> |
| --- | --- | --- | --- | --- |
| OR | GCL | Controls | 0.125 | 0.809 |
| OR | GCL | RP | 0.143 | 0.824 |
| OR | INL | Controls | 0.168 | 0.809 |
| OR | INL | RP | -0.169 | 0.809 |
| OR | IPL | Controls | 0.168 | 0.809 |
| OR | IPL | RP | 0.161 | 0.809 |
| OR | ONL | Controls | -0.1 | 0.831 |
| OR | ONL | RP | 0.249 | 0.809 |
| OR | OPL | Controls | 0.227 | 0.809 |
| OR | OPL | RP | 0.421 | 0.809 |
| OR | PhL | Controls | 0.047 | 0.912 |
| OR | PhL | RP | 0.362 | 0.809 |
| OR | RNFL | Controls | 0.071 | 0.862 |
| OR | RNFL | RP | -0.211 | 0.809 |
| OR | retina | Controls | 0.118 | 0.817 |
| OR | retina | RP | 0.37 | 0.809 |
| OT | GCL | Controls | 0.589 | 0.189 |
| OT | GCL | RP | 0.227 | 0.809 |
| OT | INL | Controls | 0.293 | 0.809 |
| OT | INL | RP | 0.009 | 0.989 |
| OT | IPL | Controls | 0.505 | 0.526 |
| OT | IPL | RP | 0.387 | 0.809 |
| OT | ONL | Controls | -0.078 | 0.846 |
| OT | ONL | RP | 0.183 | 0.809 |
| OT | OPL | Controls | 0.007 | 0.989 |
| OT | OPL | RP | 0.26 | 0.809 |
| OT | PhL | Controls | 0.003 | 0.989 |
| OT | PhL | RP | 0.244 | 0.809 |
| OT | RNFL | Controls | 0.097 | 0.832 |
| OT | RNFL | RP | -0.472 | 0.809 |
| OT | retina | Controls | 0.299 | 0.809 |
| OT | retina | RP | 0.316 | 0.809 |

Supplemental Table 2. **Correlations between foveal retinal layer thickness and white matter fibre metrics along the visual pathway in RP patients and controls.**
| Tract | Layer | Group | Pearson's r | P<sub>(FDR)</sub> |
| --- | --- | --- | --- | --- |
| OR | GCL | Controls | 0.178 | 0.809 |
| OR | GCL | RP | 0.137 | 0.831 |
| OR | INL | Controls | 0.133 | 0.809 |
| OR | INL | RP | -0.13 | 0.831 |
| OR | IPL | Controls | 0.282 | 0.809 |
| OR | IPL | RP | 0.178 | 0.809 |
| OR | ONL | Controls | -0.087 | 0.835 |
| OR | ONL | RP | 0.249 | 0.809 |
| OR | OPL | Controls | 0.358 | 0.809 |
| OR | OPL | RP | 0.304 | 0.809 |
| OR | PhL | Controls | 0.151 | 0.809 |
| OR | PhL | RP | 0.304 | 0.809 |
| OR | RNFL | Controls | 0.192 | 0.809 |
| OR | RNFL | RP | 0.006 | 0.989 |
| OR | retina | Controls | 0.216 | 0.809 |
| OR | retina | RP | 0.367 | 0.809 |
| OT | GCL | Controls | 0.443 | 0.809 |
| OT | GCL | RP | 0.425 | 0.809 |
| OT | INL | Controls | 0.152 | 0.809 |
| OT | INL | RP | -0.082 | 0.883 |
| OT | IPL | Controls | 0.616 | 0.189 |
| OT | IPL | RP | -0.248 | 0.809 |
| OT | ONL | Controls | 0.034 | 0.95 |
| OT | ONL | RP | 0.287 | 0.809 |
| OT | OPL | Controls | 0.271 | 0.809 |
| OT | OPL | RP | 0.364 | 0.809 |
| OT | PhL | Controls | 0.087 | 0.835 |
| OT | PhL | RP | 0.215 | 0.809 |
| OT | RNFL | Controls | 0.343 | 0.809 |
| OT | RNFL | RP | -0.437 | 0.809 |
| OT | retina | Controls | 0.414 | 0.809 |
| OT | retina | RP | 0.269 | 0.809 |

